# Molecular Typing of Human Respiratory Adenoviruses with Universal PCR and Sequencing Primers for Three Major Capsid Genes: Penton base, Hexon, and Fiber

**DOI:** 10.1101/2020.03.17.995373

**Authors:** Jing Zhang, Siddhartha Sridhar, Susanna K. P. Lau, June Liu, Junxian Ou, Yuqian Yan, Shan Zhao, Wendong Lan, Wenyi Guan, Jianguo Wu, Patrick C. Y. Woo, Donald Seto, Qiwei Zhang

## Abstract

Human adenoviruses (HAdVs) within species B, C, and E include the predominant types responsible for highly contagious and potentially severe respiratory infections. The traditional method to type these pathogens was based on virus neutralization and hemagglutination assays using antisera, which is both time-consuming and difficult, particularly due to the non-availability of reagents. Molecular typing based on partial characterization of HAdV hexon, as well as the restriction enzyme analysis (REA) of the genomics DNA, is difficult to identify recombinants. Here, a rapid, simple, and cost-effective molecular typing of respiratory HAdVs is presented. This incorporates three pairs of universal PCR primers that target the variable regions of the three major capsid genes, *i.e*. hexon, penton base and fiber genes. The protocol developed enables detection and typing of respiratory HAdVs within species B, C, and E, as well as of some strains within species D and F. Using this method, we surveyed a total of 100 children with acute respiratory infection caused by HAdVs in Hong Kong, Summer 2014 (July to October). 100 Throat swab specimens were collected. The samples were analyzed by PCR and the sequences were characterized by BLAST. HAdVs were detected in 98 out of 100 (98%) samples. The predominant HAdV type was species B type 3. Among the patients, 74 were of HAdV-B3 (74%), 10 were of HAdV-E4 (10%), 6 were of HAdV-C2 (21.7%), 2 were of HAdV-C6 (2%), 1 were of HAdV-B7 (2%), 1 were of HAdV-C1 (74%), and 2 were of recombinant types. The developed method allows the rapid identification of HAdVs with recombinant genomes, and bypasses the need for whole genome data, for the real-time surveillance of circulating adenovirus strains in immediate outbreaks and populations by clinical microbiologists, public health officers, and epidemiologists.

## INTRODUCTION

Human adenovirus (HAdV) is responsible for a wide spectrum of human diseases. Approximately 5-7% of respiratory illnesses diagnosed in young children are attributed to HAdVs (1, 2). Although respiratory tract infections (RTIs) caused by HAdVs are generally self-limiting, a number of severe and fatal infections have occurred in both children and adults (3-9). In addition to respiratory tract involvement, HAdVs can cause ocular, gastrointestinal, urinary bladder, and disseminated infections.

To date, more than 90 genotypes have been identified in HAdV species A-G (http://hadvwg.gmu.edu/).Among these seven species, species B types (HAdV-3, -7, -14, -16, -21, and -55) and one species E type (HAdV-4) are commonly associated with acute respiratory disease (ARD), which accounts for a high proportion of respiratory diseases in both children and adults (4, 5, 10-16).The types in species C (HAdV-1, -2, -5, -6, and -57) are generally associated with mild respiratory diseases and latent infections, while those in species D cause eye diseases and gastroenteritis. HAdV species A, F, and G are associated with gastroenteritis (17).The predominant types detected in association with disease differ among different countries or regions, and change over time (18-20).Serotypes 1 to 7 account for > 80% of HAdV infections in infants and children (21).Globally, HAdV-B3 is among the most common serotypes implicated in HAdV infections in children and adults (18, 22, 23).In South America, HAdV-B7 has been a predominant strain associated with RTI requiring hospitalization in many countries (24, 25). In Asia, HAdV-B3 and -B7 have been the predominant types associated with RTI in children (6, 12, 13, 18, 24, 26-29).In Europe, HAdV-B3 and HAdV-B7 are highly virulent and potentially lethal subtypes, especially for children (4).Historically, serotypes HAdV-B7 and -E4 predominated as a cause of ARD among military personnel in the United States (15, 20, 30, 31).

Fatal ARD outbreak caused by HAdV-3 infection was also reported (32).In immunocompromised patients in the transplant setting, some of the most commonly reported adenovirus types include HAdVC1,-C2, -C5, -A12, -A31, -B3, -B11, -B16, -B34, and -B35(33).

As a result, it is important to type HAdVs accurately and rapidly for epidemiological investigations and clinical diagnoses in order to provide information on the incidence and distribution of infections by individual types, as well as to detect and characterize novel emergent strains in the context of outbreaks (34-38).

HAdVs are traditionally typed according to serum neutralization and hemagglutination-inhibition tests, which is time-consuming. Molecular typing is based on partial characterization of HAdV hexon genes, as well as the restriction enzyme analysis (REA) of the genomics DNA, is difficult to identify recombinants. HAdV isolates with identical serum-neutralizing attribution but with distinct biological or pathogenic characteristics have been reported (39-42), challenging the traditional view of “hexon-centric” identification. With the recent development in whole genome sequencing and bioinformatics analysis, a wider range of HAdV genomes from clinical isolates have been sequenced and analyzed (43). An important finding is that recombination, scored in many cases by the three capsid genes, *i.e*., penton base, hexon, and fiber genes, and contributes to the genesis of novel and emergent pathogenic HAdVs. Among the 29 recent pathogenic genotypes identified and recognized since HAdV-52, nearly all are recombinants (39, 40, 44-52). For example, HAdV-B55 is a Trojan horse; this HAdV-B [P14H11F14] is an intertypic recombinant of HAdV-B11 and HAdV-B14 virus (46, 53, 54). It has a HAdV-14 genome chassis, including the HAdV-14 penton gene and fiber gene, but a partial HAdV-11 hexon gene, which encodes the antigenic epitopes of the virus. This virus could not only possess the virulence of HAdV-14 but also avoid the neutralizing antibody against HAdV-14. It was previously mistyped as HAdV-11a by partial hexon sequencing and REA due to incomplete gene analysis and incorrect application of the REA method (55);

Whole genome sequencing is still unfeasible for large-scale molecular epidemiological studies, as well as for rapid identification of viral pathogens during outbreaks (38, 56). Therefore, to circumvent the limitations of using only the hexon for sampling adenoviral pathogens, and also to overcome the high sequence diversity between different HAdV species, we developed a simple, rapid, cost-effective, practical, and universal typing method for the epidemiological surveillance of human respiratory adenoviruses.

At the same time, in this study, we investigated the clinical features and molecular epidemiology of HAdVs circulating among inpatient and outpatient children during the ARD outbreak in the summer of 2014 in Hong Kong, using our developed HAdV molecular typing method. This is the first report of ARD outbreak caused by HAdVs in Hong Kong as well as the first detailed molecular epidemiological survey of HAdVs in Hong Kong.

## MATERIALS AND METHODS

### Viruses and other materials

Adenovirus strains HAdV-B3, -B7, -B11, -B14, -B21, -B55, -C5, -D19, –E4, and -F41 have been isolated, identified, and archived in our laboratory (6, 10, 13, 46, 58, 59). Taq PCR Master Mix kits (Takara; Japan), QIAamp DNA Mini Kit (QIAgen; China), PCR cleanup kit (Axygen) were used according to manufacturer instructions. DL10000 and DL2000 DNA Markers were the product of TAKARA Corp.

### Patients and clinical specimens

From July 2014 to October 2014, there was a sudden increase in pediatric outpatients and inpatients with influenza-like symptoms in Queen Mary Hospital, Hong Kong. nasopharyngeal swab specimens were collected from patients and respiratory viral pathogens were detected.

This study protocol was approved by the Institutional Ethics Committee of Mary Hospital and was carried out in accordance with the approved guidelines. Legal guardians of all underage participants gave signed informed consent for their participation in this study. Data records of the samples and sample collection were de-identified and anonymized.

Detection of human adenovirus and other common respiratory pathogens with Immunofluorescent Kits

Total nucleic acid was extracted from respiratory specimens using a QIAamp DNA Mini Kit (QIAgen), in accordance with the manufacturer’s protocol. Adenovirus and other ten common respiratory pathogens were detected using Taqman real-time PCR kit according to the manufacturer’s protocol, including influenza A virus (infA), influenza B virus (infB), parainfluenza virus (PIV), respiratory syncytial virus (RSV), enterovirus (EV), human metapneumovirus (hMPV), bocavirus (BOV), rhinovirus (RHV), Mycoplasma pneumoniae (MP), and Chlamydia pneumonia (CP) (Co. LTD;China).

### Adenovirus culture and isolation

The 100 HAdV-positive throat swab specimens collected from 100 patients were inoculated onto A549 cells and were cultured in a maintenance medium (Minimal Essential Medium containing 2% fetal bovine serum, 100 U/ml penicillin G and 100 μg/ml streptomycin) at 37°C in an atmosphere containing 5% (v/v) carbon dioxide, and cytopathic effect (CPE) was monitored for 5-7 days, If no CPE was observed, the cells would be freezed and thawed for three times and passaged in A549 again to check CPE.

### PCR primers design and PCR amplification

The penton base, hexon, and fiber gene sequences from HAdV-A18, -B3, -B7, -B11, -B14, -B16, -B21, -B34, -B35, -B50, -B55, -B66, -B68, -C1, -C2, -C5, -C6, -D19, -D37, -E4, -F41, and -G52 were obtained from GenBank, and were aligned using ClustalW to find regions with high sequence similarities. Primers targeting the three capsid genes were designed based on these bracketing conserved regions and ordered from Invitrogen (Guangzhou).

PCR reactions were conducted in a total volume of 20 μL containing 1× Taq Master Mix (10 μL), primer F (10 μmol/L, 0.5 μL), primer R (10μmol/L, 0.5 μL), DNA template (1 μL), and water (8 μL). Primers Penton-F and Penton-R were designed for both PCR amplifying and DNA sequencing of the penton base gene. PCR was performed as follows: 94 °C for 1 min; 34 cycles of 94 °C for 30 s; 52 °C for 30 s; and 72 °C for 100 s, with a final extension of 72 °C of 10 min. Primers HVR-F and HVR-R were used for the hexon gene. PCR was performed as follows: 94 °C for 1 min; 34 cycles of 94 °C for 30 s; 52°C for 30 s; and 72 °C for 100 s, with a final extension of 72 °C of 10 min. Primers Fiber-F and Fiber-R were designed for the fiber gene. The PCR amplification conditions for this are as follows: 94 °C for 1 min; 34 cycles of 94 °C for 30 s; 52 °C for 30 s; and 72 °C for 72 s, with a final extension of 72 °C of 10 min.

### Sequencing and molecular typing of clinical adenovirus specimens

All three major capsid genes of the HAdV isolates were PCR-amplified by the corresponding pairs of universal primers. These PCR products were purified and sequenced directly with both PCR primers by Invitrogen (Guangzhou; China). The assembled DNA contigs were characterized by a BLAST survey of the NCBI GenBank database. The molecular type of each clinical specimen was determined in accordance to the highest sequence identities.

### Genome sequences used for alignments

Archived genome sequences from GenBank were used for the alignments of the penton base, hexon, and fiber genes. Their accession numbers are as follow: HAdV-A12 (X73487), HAdV-B3 (DQ099432), HAdV-B3 (AY599834), HAdV-B7 (AY594255), HAdV-B7 (KC440171), HAdV-B11 (AY163756), HAdV-B14 (AY803294), HAdV-B16 (AY601636), HAdV-B21 (AY601633), HAdV-B34 (AY737797), HAdV-B35 (AY128640), HAdV-B50 (AY737798), HAdV-B55 (JX491639), HAdV-C1 (AC_000017), HAdV-C2 (J01917), HAdV-C5 (AC_000008), HAdV-C6 (KY268129), HAdV-D9 (AJ854486), HAdV-E4 (AY594253), HAdV-F40 (KU162869), HAdV-G52 (DQ923122), SAdV-25 (AC_000001), SAdV-26 (FJ025923).

### Phylogenetic analysis

The Molecular Evolutionary Genetics Analysis (MEGA) version 7.0 software (www.megasoftware.net/) was used for phylogenetic analyses of penton base, hexon, and fiber genes from the clinical specimens, along with additional reference sequences retrieved from GenBank database. Phylogenetic trees were constructed using the maximum parsimony method with a bootstrap test of 1000 replicates and the Tree-Bisection-Reconnection (TBR) model.

## RESULTS

### Three pairs of universal primers targeting the three major capsid genes: penton base, hexon, and fiber genes

Three pairs of universal primers were designed for PCR amplification and sequencing of HAdV penton base, hexon, and fiber genes. Primers Penton-F and Penton-R were designed based on the conserved regions of the penton base sequences. Within the alignment of sequences, primer sequences for Penton-F and Penton-R are highly conserved in the majority of penton base genes, which ensures that all of the analyzed adenovirus types within species A to G could be PCR-identified (Fig. 1(B)). The resultant PCR product is 1253bp (Table 1), located within penton base gene and contains the variable region HVR1 and the RGD loop (Fig. 1(A)).

**Table 1.**
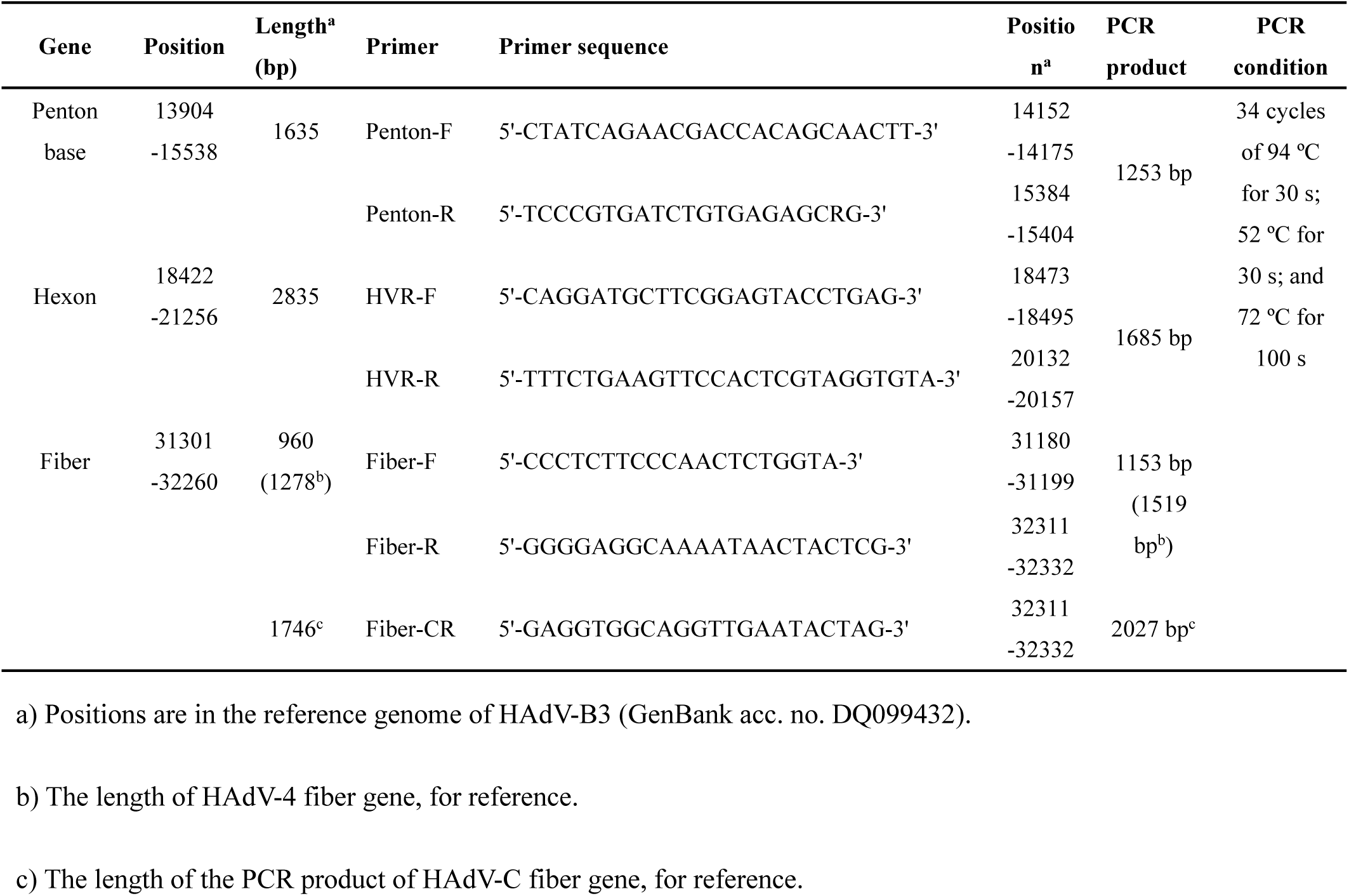
Universal primers for the detection, typing and sequencing of human adenoviruses. Sequences, genome locations, and the resultant predicted PCR product sizes are noted for each of the three major capsid protein genes.

**Fig. 1.**
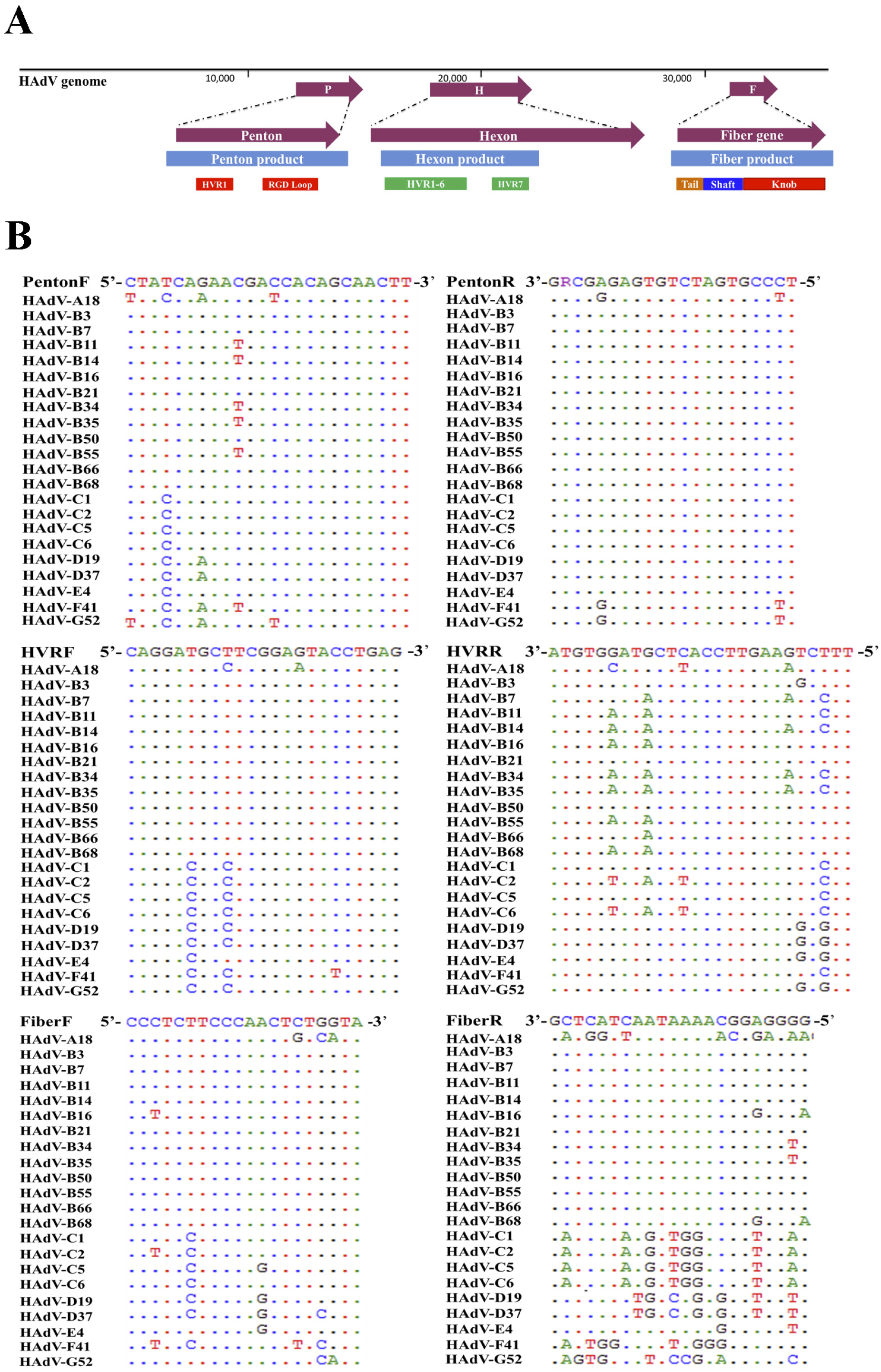
(A). Alignment of human adenovirus (HAdV) universal primers for HAdV-A18, B3, -B7, B-11, -B14, -B16, -B21, -B34, -B35, -B50, –B55, -B66, -B68, -C1, -C2 –C5, -C6, -D9, -D19, -D37, -E4, -F41, and -G52. Divergence from the primer sequences are shown for the isolates tested. Each virus is identified with type number as well as species demarcation for reference. Dots represent identical bases, base changes are noted, and the bases are color-coded for visual comparisons. And (B). **Schematic adenovirus genome, with the primer positions and the resultant PCR products noted for the three major capsid protein genes: penton base (left), hexon (middle), and fiber (right) genes**. The relative locations of the capsid protein genes in the HAdV genome are noted. These genes contain the typing and molecular characterization information for HAdVs. The purple arrows indicate the genes and their locations; the blue bars indicate the PCR products and the relative length; and the other colored bars indicate the important domains within each gene, including hypervariable regions (HVRs and RGD Loops), as well as the tail, shaft, and knob domains.

For the hexon, the universal amplification primers HVR-F and HVR-R were designed. The primer sequences are conserved in HAdV types across all of the species analyzed, yielding a PCR product of about 1685 bp (Table 1).This amplicon contains the seven hypervariable regions (HVRs) comprising Loops 1 and 2 (Fig. 1(A)).

Primers Fiber-F and Fiber-R were designed for fiber gene. Point mutations of primer Fiber-F exist only in one or two nucleotides located in the middle of the primers; this ensures an effective PCR amplification of the different HAdV types (Fig. 1(A)). These primers amplify HAdV types from species B, D, and E, yielding a PCR product of about 1153 bp for HAdV-B and -D, and 1519 bp for HAdV-E4. However, during the high variation and long fiber gene (about 1746 bp) of species C, Fiber-R matches poorly with types in HAdV-C. To compensate, another primer Fiber-CR was designed to completely match the sequences within species C (Table 1).

### PCR identification of HAdV-B3, -B7, -B11, -B14, -B21, -B55, -C5, -D19, -E4, and -F41 using universal primers targeting the three capsid genes

Genomic DNA of strains of HAdV-B3, -B7, -B11, -B14, -B21, -B55, -C5, -D19, -E4, and -F41 were extracted and detected by PCR using these three pairs of universal primers. All of these HAdV types are detected by PCR amplification. The PCR products were specific, yielding single distinct products. The sizes of the PCR products are identical with the expected sizes: 1.2 kb (penton base), 1.6 kb (hexon), and 1.1 kb (fiber), respectively. There are two exceptions: one is the HAdV-E4 fiber gene (1519 bp) and the other is the HAdV-C5 fiber gene (2027bp). Both gene products are longer than their counterparts in the other HAdV types.

### Demographic characteristics and clinical features of HAdV-positive children with ARD in Hong Kong

Total 100 HAdV-positive samples were collected and analyzed, (Table2). It showed that the male and female rates were 50% and 49%, respectively. No significant gender difference was found. The median age with HAdV infection was 4 years, ranging from 0.5 year to 14 years old, of which, 60% were under 5 years. Of the 100 cases, 93 (93%) were hospitalized; the median duration of hospitalization was 3 days (1-36 days). Fourteen HAdV-positive cases were co-infected with EV/RV, 2 with RSV, 1 with influenza C.

The clinical diagnoses included one case of fatal pneumonia caused by HAdV-3, 3 cases of bronchitis, 5 cases of diarrhea, 2 cases of febrile seizure, 2 cases of rash, and 74 cases of upper respiratory tract infection (URTI) (Table 2).

**Table 2.** Comparison of demographic and clinical characteristics of 100 children with ARD according to HAdV type in Hong Kong, summer 2014.

| HAdV species |  | All | HAdV-B3 | HAdV-C1 | HAdV-C2 | HAdV-E4 | HAdV-C6 | HAdV-B7 | Recombinant<br>Coinfect |
| --- | --- | --- | --- | --- | --- | --- | --- | --- | --- |
| Numbers |  | 100 | 74 | 1 | 3 | 10 | 2 | 1 | 7 |
| Gender (M/F |  | 50/49 | 38/35 | 0/1 | 2/1 | 3/7 | 1/1 | 1/0 | 5/2 |
| Age (range) |  | 4 (0.5-14) | 2.5 (1-14) | 4 (4) | 3 (0.5-7) | 5 (1-11) | 3.5 (3-4) | 7 (7) | 2 (1-9) |
| Nationality | Chinese | 86 | 63 | 1 | 1 | 9 | 2 | 1 | 7 |
|  | Occidental | 2 | 2 | - | - | - | - | - | - |
|  | Korean | 1 | 1 | - | - | - | - | - | - |
|  | Indian | 1 | 1 | - | - | - | - | - | - |
|  | Nepalese | 3 | 1 | - | 2 | - | - | - | - |
| Hospitalized Numbers |  | 93 | 68 | 1 | 3 | 9 | 2 | 2 | 6 |
| Hospitalized day (range) |  | 3 (1-36) | 3 (1-36) | 3 (3) | 3 (2-4) | 2 (1-15) | 2.5 (2-3) | 4 (4) | 2 (2-3) |
| Co-infection | EV/RV | 14 | 10 | - | 1 | 2 | - | - | - |
|  | RSV | 2 | 1 | - | - | 1 | - | - | - |
|  | Flu C | 1 | 1 | - | - | - | - | - | - |
| Clinical Diagnosis |  |  |  |  |  |  |  |  |  |
| URTI |  | 74 | 55 |  | 3 | 8 | 1 |  | 6 |
| Diarrhea |  | 5 | 3 | 1 |  |  |  |  | 1 |
| Bronchiolitis |  | 3 | 2 |  |  |  |  |  |  |
| Febrile seizure |  | 3 | 3 |  |  |  |  |  |  |
| Pneumonia* |  | 1 | 1* |  |  |  |  |  |  |
| Rash |  | 2 | 2 |  |  |  |  |  |  |
| Adenoviremia transplant recipient |  | 1 | - |  |  | 1 |  |  |  |
| Conjunctivitis/ Febrile seizure |  | 1 | 1 |  |  |  |  |  |  |
| Fever/ GE |  | 1 | - |  |  |  |  | 1 |  |
| Intussusception |  | 1 | - |  |  |  | 1 |  |  |
| N/A |  | 8 | 7 |  |  | 1 |  |  |  |

**\*: dead.**

### Molecular typing of HAdV clinical samples

One hundred throat swab specimens were subsequently detected by PCR using the three pairs of universal primers. The PCR products were sequenced. BLAST analysis confirmed that 74 cases were HAdV-B3, 10 cases were HAdV-E4, 6 cases were HAdV-C2, 2 cases were HAdV-C6, 1 case were HAdV-B7, 1 case was HAdV-C1, and 4 cases were infected by recombination HAdVs or co-infected by different types of HAdVs; the other 2 cases were PCR negative (Fig. 2A). The 4 cases infected by recombinants or co-infected by different HAdV types were HK39 [P7H55F7], HK42 [P1H1F3], HK61 [P1H5F2], and HK76 [P1H2F2].

**Fig.2.**
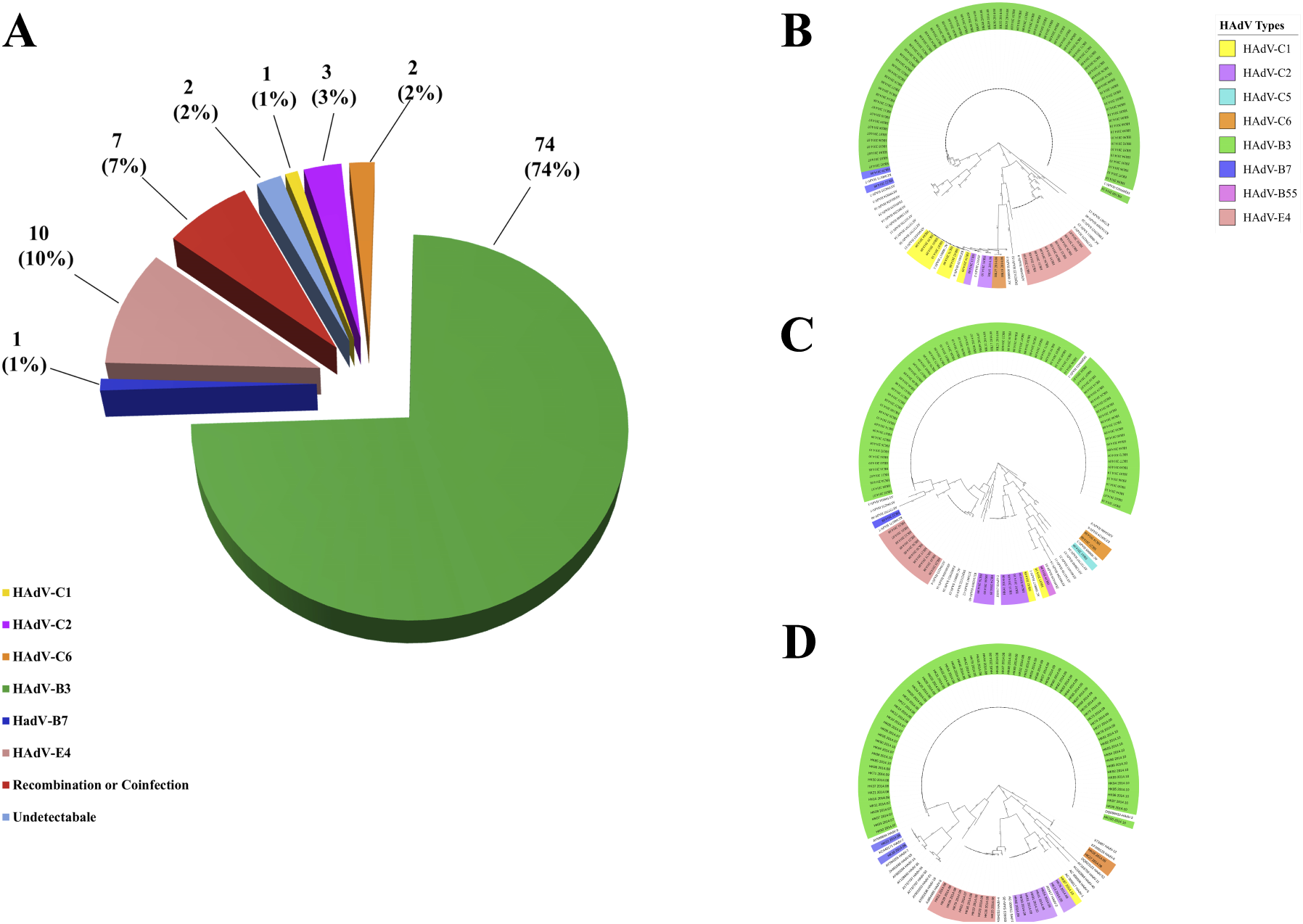
A. HAdV type distribution in hospitalized children with ARIs from July through October, 2014. And Panels (B), (C), and (D) display the phylogenetic relationships of the penton base, hexon, and fiber genes, respectively, of the 98 clinical HAdV strains. Additional reference sequences were retrieved from GenBank to provide context. The phylogenetic trees were generated based on the Tree-Bisection-Reconnection (TBR) model by MEGA 7.0 (www.megasoftware.net/) using the maximum parsimony method with 1,000 boot-strap replicates and default parameters. The percentage of trees in which the associated taxa clustered together is shown next to the branches. The scale bar is in units of nucleotide substitutions per site.

### Phylogenetic analysis of HAdV clinical samples

Phylogenetic analysis of the hexon, penton base, and fiber gene of Hong Kong isolates were shown in fig 2.The phylogenetic analysis results were consistent with the BLAST results. Panels (B), (C), and (D) display the phylogenetic relationships of the penton base, hexon, and fiber genes, respectively, of the 100 clinical HAdV strains. HAdV-B3 (n = 74) was the most prevalent types. Three capsid genes formed a subclade with another China HAdV-3 isolate Guangzhou01 circulating in 2004(10).HAdV-E4 was the second mostprevalent strain (n = 10).

### GenBank accession numbers of the capsid protein genes from clinical isolates

The PCR products of the penton base, hexon, and fiber genes from the thirty throat swab specimens were sequenced. These were submitted to GenBank. GenBank accession numbers are as follows:

## DISCUSSION

PCR and sequencing analysis of microbial DNA have been used for quick identification and better characterization of the pathogens (60). The identification of HAdV types was traditionally performed by hexon protein analysis, *e.g*., virus neutralization, which depends on the interaction between type-specific antisera and type-specific antigenic epitopes in HVRs of the hexon protein (61), or by limited hexon gene sequencing (62). The time-consuming and labor-intensive epitope detection method is seldom used today; the limited partial hexon sequencing provides partial and incomplete identification, as any recombinant HAdV will be missed. As a gold standard, whole genome sequencing and analysis is the most accurate method to identify, characterize, and type HAdVs. This is borne out by recent recognition of new genotypes that include recombinant HAdVs identified by this whole genome analysis method, including important pathogenic and emergent HAdVs. As an example, HAdV-D53 was recognized as a new genotype because genome recombination was detected amongst the three major capsid genes. The penton base, hexon, and fiber genes originated from HAdV-D37, HAdV-D22, and HAdV-D8, respectively (51). Another example is the emergent genotype HAdV-B55 (40, 56). This is a highly contagious recombinant respiratory pathogen that contains the hexon epitope from a urinary tract pathogen HAdV-B11 and the penton base and fiber genes from a respiratory tract pathogen HAdV-B14.

Given that whole genome sequencing is still relatively cost-prohibitive, particularly for large numbers of samples comprising outbreak and population sampling projects, and that genome recombination may only be indicated by assaying marker genes across the genome, such as the penton base, hexon, and fiber genes, a simple, rapid, cost-effective, practical, and universal detection and typing method for characterizing HAdVs is presented in this study. This protocol calls for using three pairs of universal PCR primers to target variable regions of the three capsid genes to provide products for characterizing the adenoviral isolates. The subsequent amplicon sequencing and BLAST analysis provides information as to the type identity and whether there is any recombination across the genome. This method was validated by typing 30 clinical specimens successfully. In practice, all three pairs of universal primers that we chose and optimized work for HAdV-B3, -B7, -B11, -B14, -B21, -B55, -C5, -D19, and -E4. Because the HAdV-C fiber gene sequences are phylogenetically distinct from the other species, to compensate, we designed a specific primer Fiber-CR, which worked well for the HAdV-C fiber amplification. The universal primers for penton base and hexon genes successfully amplified the assayed types across species B to F, although the genomic sequences between different species are diverse. Isolates from the set of genotypes representing species HAdV-B, -C, -D, and -E can be detected and type-identified by this protocol with these universal primers, including putative recombinants: the HAdV-B55 isolates.

In contrast to a commonly used HAdV typing protocol published by Lu *et al*. (62) and others which were based solely on the PCR amplification and/or sequencing of the HAdV hexon gene (56, 63-65), the method presented in this report is more informative. Additionally, the Lu *et al*. protocol targets only the HVR1-6 (62) rather than the entire epitope. Although HVR1-6 contain type-specific epitopes, the adjacent HVR-7 region does as well (65, 66). And as noted, these hexon-centric methods do not identify recombination across the genome, which appears to be an important molecular evolution mechanism in the genesis of novel HAdVs, as noted by the recent characterization and recognition of several emergent human adenoviral pathogens (39, 40, 44-52). On the contrary, our PCR amplification and DNA sequencing method not only targets three genes that essentially span the genome, but includes seven HVRs, which will identify any recombination within the hexon epitopes as well. The three PCR reactions can be performed concurrently and rapidly, saving the detection time. This method will economically provide the identification and characterization of HAdVs, particularly recombinants, in the real-time surveillance, sampling, and screening of circulating large numbers of adenovirus isolates during outbreaks and in populations for clinical microbiologists, public health officers, and epidemiologists.

## ACKNOWLEDGEMENTS

We thank the Guangzhou Women and Children’s Medical Center, Zhujiang Hospital, Hubei CDC, and Guangdong CDC for providing clinical throat swab specimens.

## FUNDING INFORMATION

This work was supported by grants from the National Key Research and Development Program of China (2018YFE0204503), National Natural Science Foundation of China (81730061), Natural Science Foundation of Guangdong Province (2018B030312010) as well as the Guangzhou Healthcare Collaborative Innovation Major Project (201803040004 and 201803040007).

